# Charcot-Leyden Crystals activate the NLRP3 inflammasome and cause IL-1β inflammation

**DOI:** 10.1101/252957

**Authors:** Juan Francisco Rodríguez-Alcázar, Marco Antonio Ataide, Gudrun Engels, Christine Schmitt-Mabmunyo, Natalio Garbi, Wolfgang Kastenmüller, Eicke Latz, Bernardo S Franklin

## Abstract

Charcot-Leyden crystals (CLCs) are Galectin-10 protein crystals that can form after eosinophils degranule. CLCs can appear and persist in tissues from patients with eosinophilic disorders, such as asthma, allergic reactions, fungal, and helminthic infections. Despite abundant reports of their occurrence in human disease, the inflammatory potential of CLCs has remained unknown. Here we show that CLCs induce IL-1β release upon their uptake by primary human macrophages *in vitro,* and that they induce inflammation *in vivo* in mouse models of acute peritonitis and bronchitis. CLC-induced IL-1β was dependent on NLRP3 and caspase-1, and their instillation in inflammasome reporter mice promoted the assembly of ASC complexes and IL-1β secretion in the lungs. Our findings reveal that CLCs are recognized by the NLRP3 inflammasome, which may sustain inflammation that follows eosinophilic inflammatory processes.

## INTRODUCTION

Modifications in the physicochemical properties of certain proteins, lipids, or metabolites can result in phase transition leading to crystal formation or non-physiological aggregation (Franklin et al., 2016; Mulay and Anders, 2016). Crystals can arise in tissues when endogenous material, such as minerals, cholesterol, and uric acid deposit. They can also be introduced from exogenous sources by inhalation (silica, asbestos, airborne particulate matter), or via injections (medical nanomaterials, vaccine adjuvants) (Franklin et al., 2016). Crystalline or aggregated materials are sensed by innate immune cells, which are equipped with germ-line encoded pattern recognition signaling receptors (PRRs). Innate immune cells survey tissues and remove accumulated extracellular material, pathogens, and residues of cellular demise. Sensing of crystals induces potent inflammatory responses, a property that has been exploited for adjuvanticity in vaccine formulations. Whereas several membrane-bound PRRs are involved in the recognition and uptake of crystals (Tsugita et al., 2017; Sheedy et al., 2013), the cytosolic PRR NLRP3 senses excessively phagocytosed crystals. NLRP3 activation causes the formation of an inflammasome that leads to activation and release of pro-inflammatory cytokine interleukin-1 beta (IL-1β) (Hornung et al., 2008; Duewell et al., 2010; Masters et al., 2010; Halle et al., 2008; Martinon et al., 2006b; Dostert et al., 2008). IL-1β plays important roles in crystal-driven inflammatory diseases and, together with NLRP3, represents a promising therapeutic target for a range of crystal-mediated diseases (Franklin et al., 2016; Mulay and Anders, 2016).

Inflammation occurs not only as a consequence of crystal buildup, but it can itself promote crystallization. During inflammation immune cells are recruited to the affected tissue. Among them, eosinophils and basophils play important roles in allergic reactions, fungal and helminthic infections, and in the regulation of a variety of autoimmune disorders. These cells can expel highly cytotoxic proteins from their secretory granules, which mediate the marked physiological changes associated with eosinophil and basophil inflammation. A predominant protein present in eosinophilic and basophilic granules is Galectin-10 (LGASL10), also known as Charcot-leyden protein (Golightly et al., 1992; Archer and Blackwood, 1965). Galectin-10 can cluster and form Charcot-Leyden Crystals (CLCs), which are colorless elongated hexagonal bipyramidal structures that can reach up to 50 μm in length. Since their first report, more than 160 years ago, CLCs have been shown to spontaneously form *in vitro* from lysates of eosinophils (Weller et al., 1980) and basophils (Ackerman et al., 1982), and have been found to accumulate in tissues from patients with asthma (Dor et al., 1984), fungal allergic reactions (el-Hashimi, 1971; Katzenstein et al., 1983; DeShazo et al., 1997), helminthic infections (Kaplan et al., 2001), and myeloid leukemia (Nashiro et al., 2016; van de Kerkhof et al., 2015; Manny and Ellis, 2012). Despite abundant reports showing the appearance of CLCs in tissues from patients with eosinophilic disorders, these crystals are still regarded as inert remains of eosinophil activity, with unrecognized immune function. Advances in this field have been hindered by the lack of a suitable animal model to study CLC formation *in vivo*. Mice lack the gene *LGASL10*, which encodes for Galectin-10. Therefore, it remains to be demonstrated whether CLCs are inflammatory *in vitro* and *in vivo*.

Here we show that CLCs are promptly phagocytosed *in vitro* by human macrophages, and this leads to the release of the pro-inflammatory cytokine IL-1β. CLC-induced IL-1β in macrophages occurred through the activation of caspase-1 following the assembly of the NLRP3 inflammasome. Furthermore, we show that CLCs promoted inflammation *in vivo*, characterized by neutrophil infiltration, phagocytosis of the crystals, and IL-1β release. Additionally, CLCs induced the assembly of inflammasomes (ASC specks) *in vivo* after their instillation into the lungs of ASC-mCitrine transgenic reporter mice.

Our findings uncover immune stimulatory features of CLCs and shed new light into the likely consequences caused by inflammatory sequelae that follow eosinophil infiltration in tissues.

## RESULTS AND DISCUSSION

### Production and characterization of Charcot-Leyden crystals

To investigate the inflammatory potential of CLCs, we first produced CLCs from whole cell lysates of AML14.3D10 cells, a subclone of the AML14 cell line that was established from a 68-year-old patient with FAB M2 acute myeloid leukemia (Baumann and Paul, 1998). Both the parental AML14 cell line and the AML14.3D10 subclone are established tools for the study of eosinophil biology (Paul et al., 1994). We produced CLCs *in vitro* by incubating pre-cleared lysates of AML14.3D10 cells at 4 °C for 16 hours (Fig. 1a, and Fig. S1a) as previously described (Ackerman et al., 1980). The resulting crystals displayed the hexagonal bipyramidal morphology characteristic of CLCs (Fig. 1b). Coomassie Brilliant Blue staining of a gel loaded with SDS-denatured crystals revealed that CLC preparations contained mainly one protein (Fig. 1c, and Fig. S1a), and immunoblotting with specific anti-Galecin-10 mAb showed that the CLC preparations were composed of Galectin-10 (Fig. 1d and Fig. S1a).

**Fig. 1.**
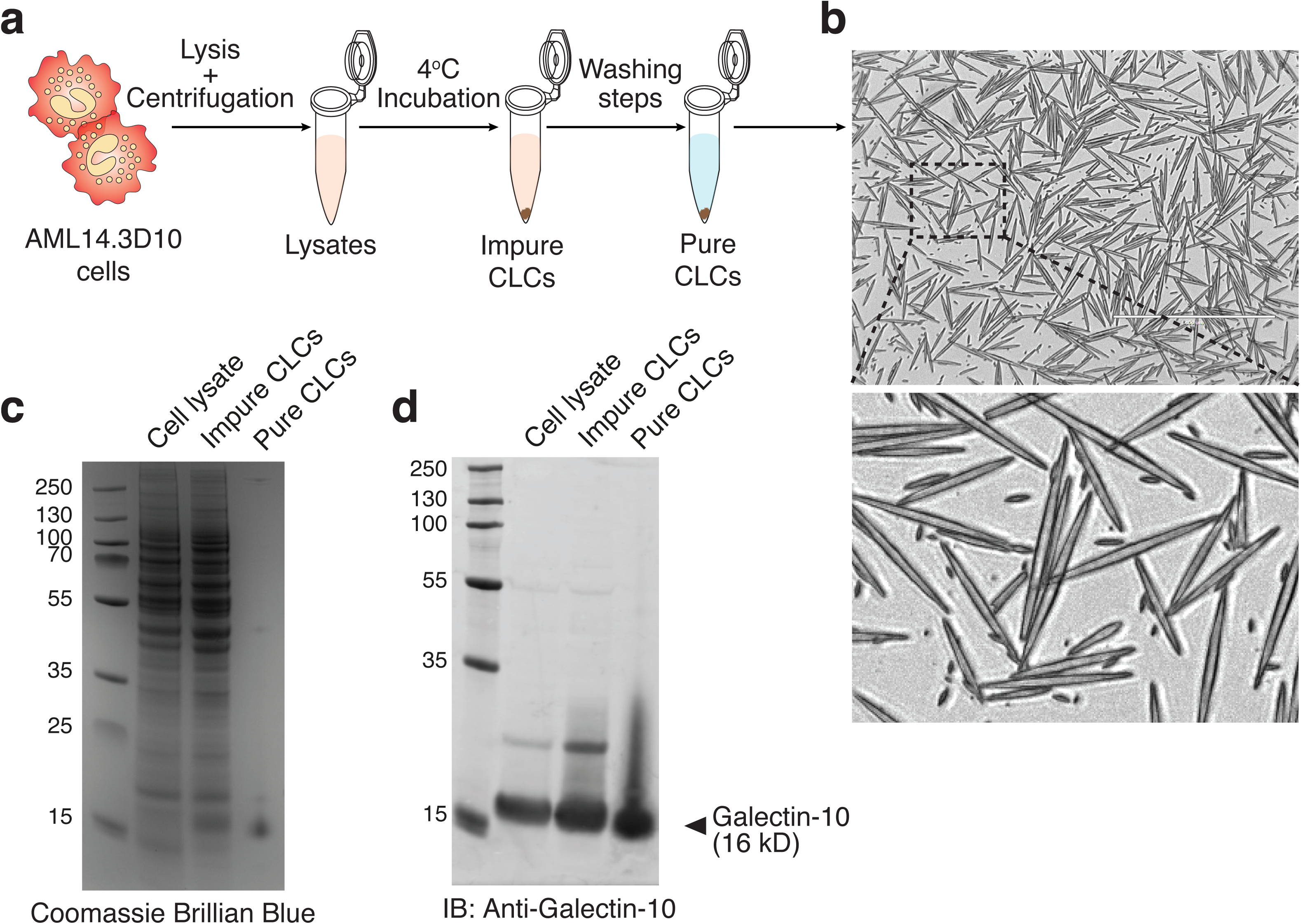
Production and characterization of CLCs. (**a**) Schematics of the generation of CLCs from AML14.3D10 cells. (**b**) Wide-field imaging of a representative pure CLC fraction generated as shown in **a**. Scale bars: 400 µm. Insert is a 20X magnification of the area outlined at top. (**c**) Coomassie-stained protein gel and (**d**) immunoblot for Galectin-10 of representative fractions of lysates of AML14.3D10 cells, impure CLCs, and pure CLCs that were generated as shown in **a**. Data is representative of two independent experiments.

### Phagocytosis of CLCs leads to IL-1β secretion by human macrophages

We next generated primary human monocyte-derived macrophages (hMDMs) differentiated from CD14^+^ monocytes isolated from the peripheral blood mononuclear cells (PBMCs) from buffy coats of healthy donors. These cells were primed with LPS and exposed to CLCs and their phagocytic potential against the CLCs was assessed by microscopy. We noticed that hMDMs promptly engulfed CLCs of variable sizes (Fig. 2a, and Movie S1).

**Fig. 2.**
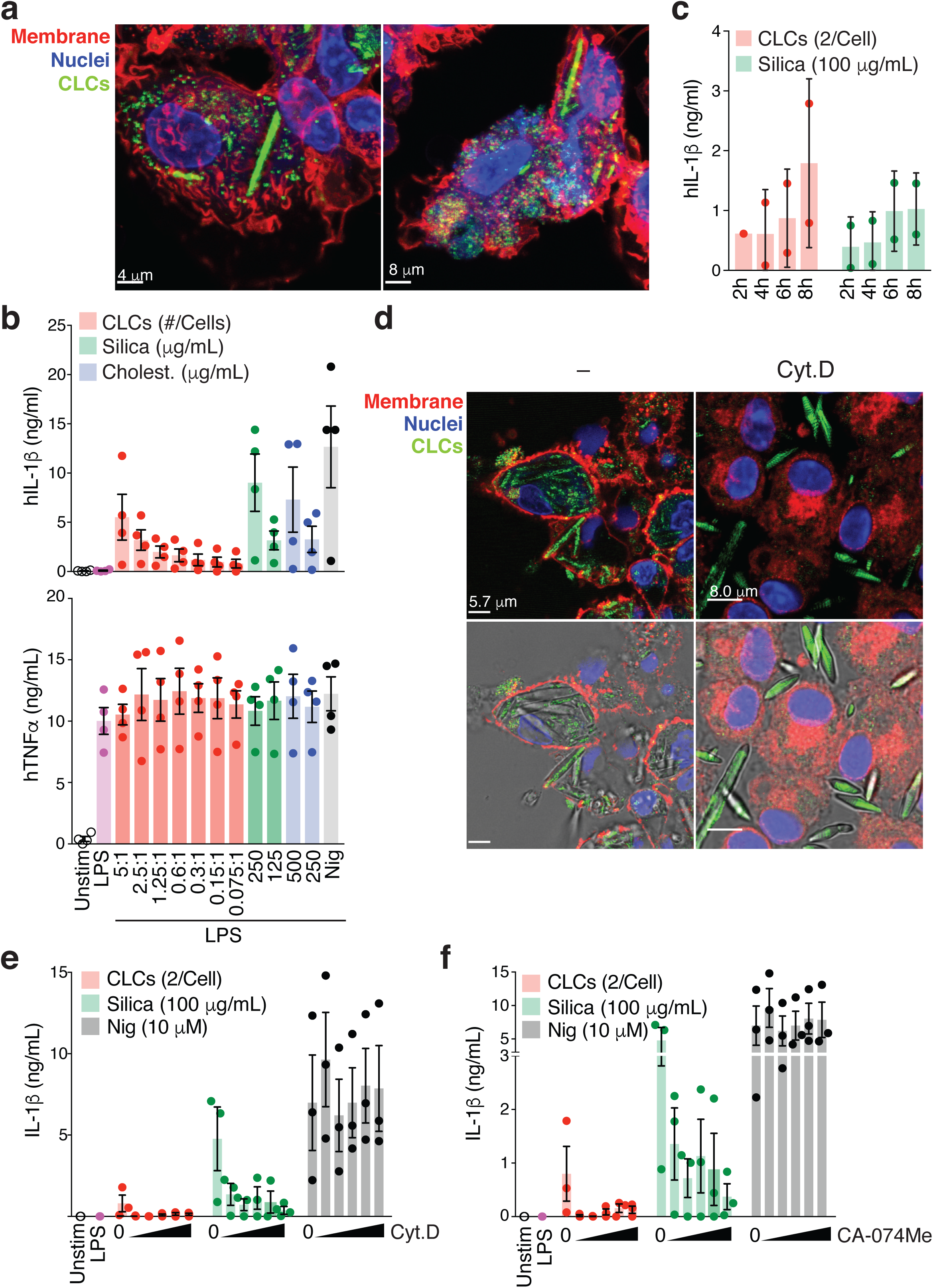
Phagocytosis of Charcot-Leyden Crystals (CLCs) causes IL-1β secretion by human macrophages *in vitro*. (**a**) Confocal imaging of LPS-primed (2 ng/mL, for 3 hours) primary human macrophages (hMDMs) stimulated with purified CLCs for 6 hours. Plasma membrane (red, WGA-AF555), Nuclei (blue, DRAQ5), and CLCs (green, laser reflection). Scale bars (4 µm left panel, and 8 μm right panel). Data is from one representative out of three independent experiments. (**b**) HTRF measurement of IL-1β and TNFα from the supernatants of hMDMs that were either left untreated, primed with LPS, or primed with LPS and stimulated for 6 hours with the indicated ratios of CLCs/macrophage, or the indicated concentrations of silica crystals, cholesterol crystals, or for 1.5 hours with nigericin (Nig). Each symbol represents the values from hMDMs generated from different donors. Bars represent mean and SEM of pooled data from four independent experiments, with different CLC preparations. (**c**) HTRF measurement of IL-1β in hMDMs primed as in **a** and stimulated with CLCs (2 per cell) or silica crystals (100 μg/mL) for the indicated time points. Bars represent mean and SD of pooled data from two independent experiments. Each symbol represents the values from a different donor. (**d**) Confocal imaging of LPS-primed hMDMs that were pre-treated or not with 5 μΜ of cytochalasin D, 30 minutes before incubation with CLCs (2 per cell). (**e**) HTRF measurement of IL-1β from the supernatants of LPS-primed hMDMs that were pre-treated with increasing concentrations of cytochalasin D (0, 0.3, 0.6, 1.25, 2.5 or 5 μM). Bars represent mean and SEM of pooled data from three independent experiments. Each symbol represents the values from different donors. (**f**) HTRF measurement of IL-1β from the supernatants of LPS-primed hMDMs that were pre-treated with increasing concentrations of the cathepsin-B inhibitor CA-074 Me (0, 1.25, 2.5, 5, 10, 15 μM). Bars represent mean and SEM of pooled data from three independent experiments. Each symbol represents the values from different donors.

The pro-inflammatory cytokine IL-1β is a key driver of crystal-induced inflammation in several chronic inflammatory diseases (Hornung et al., 2008; Duewell et al., 2010; Masters et al., 2010; Halle et al., 2008; Martinon et al., 2006b; Dostert et al., 2008). We thus investigated whether phagocytosis of CLCs resulted in IL-1β release by hMDMs. We found that, similar to the well-established inflammasome activators silica and cholesterol crystals, exposure of LPS-primed hMDMs to CLCs resulted in a dose– and time-dependent release of IL-1β from these cells (Fig. 2b-c). Similar to reports using other crystals, LPS priming was required for the release of IL-1β by *in vitro* CLC-stimulated hMDMs (Table S1). Importantly, phagocytosis was required for CLCs-induced IL-1β release by hMDMs *in vitro*, as treatment of cells with the actin polymerization inhibitor cytochalasin D impaired their phagocytic activity and diminished their capacity to release IL-1β in response to CLCs stimulation (Fig. 2d-e). As expected, cytochalasin D inhibited the IL-1β release mediated by silica crystals, while the response to nigericin, a soluble inflammasome activator that acts as an antiporter of H^+^ and K^+^, remained unaffected.

### CLCs-induced IL-1β release *in vitro* depends on NLRP3 activation

Phagocytosis of crystalline material and insoluble protein aggregates can ultimately result in lysosomal damage and the release of lysosomal content into the cell cytosol. Lysosomal damage can activate NLRP3, which then recruits the adapter protein ASC and caspase-1, a protein complex that enables maturation of pro-IL-1β by caspase-1-mediated proteolysis (Hornung et al., 2008). To investigate whether lysosomal damage is involved in CLCs-induced inflammasome activation, we pre-treated hMDMs with CA-074 Me, a compound that inhibits lysosomal damage-mediated NLRP3 activation (Hornung et al., 2008). Indeed, CA-074 Me inhibition blocked CLC-induced IL-1β release in a dose-dependent manner (Fig. 2f), suggesting that lysosomal damage is a likely mechanism by which CLCs can induce IL-1β secretion by macrophages. Inflammasome assembly can be visualized by ASC transition from a diffused cytosolic localization into a dot-like structure called the ASC speck (Stutz et al., 2013; Hoss et al., 2017). We therefore performed immunofluorescence staining of ASC in LPS-primed hMDMs stimulated with CLCs. Using fluorochrome-conjugated anti-ASC antibodies, or isotype matched fluorochrome-conjugated IgGs, we revealed the formation of ASC specks in hMDMs that phagocytosed CLCs *in vitro* (Fig. 3a).

**Fig. 3.**
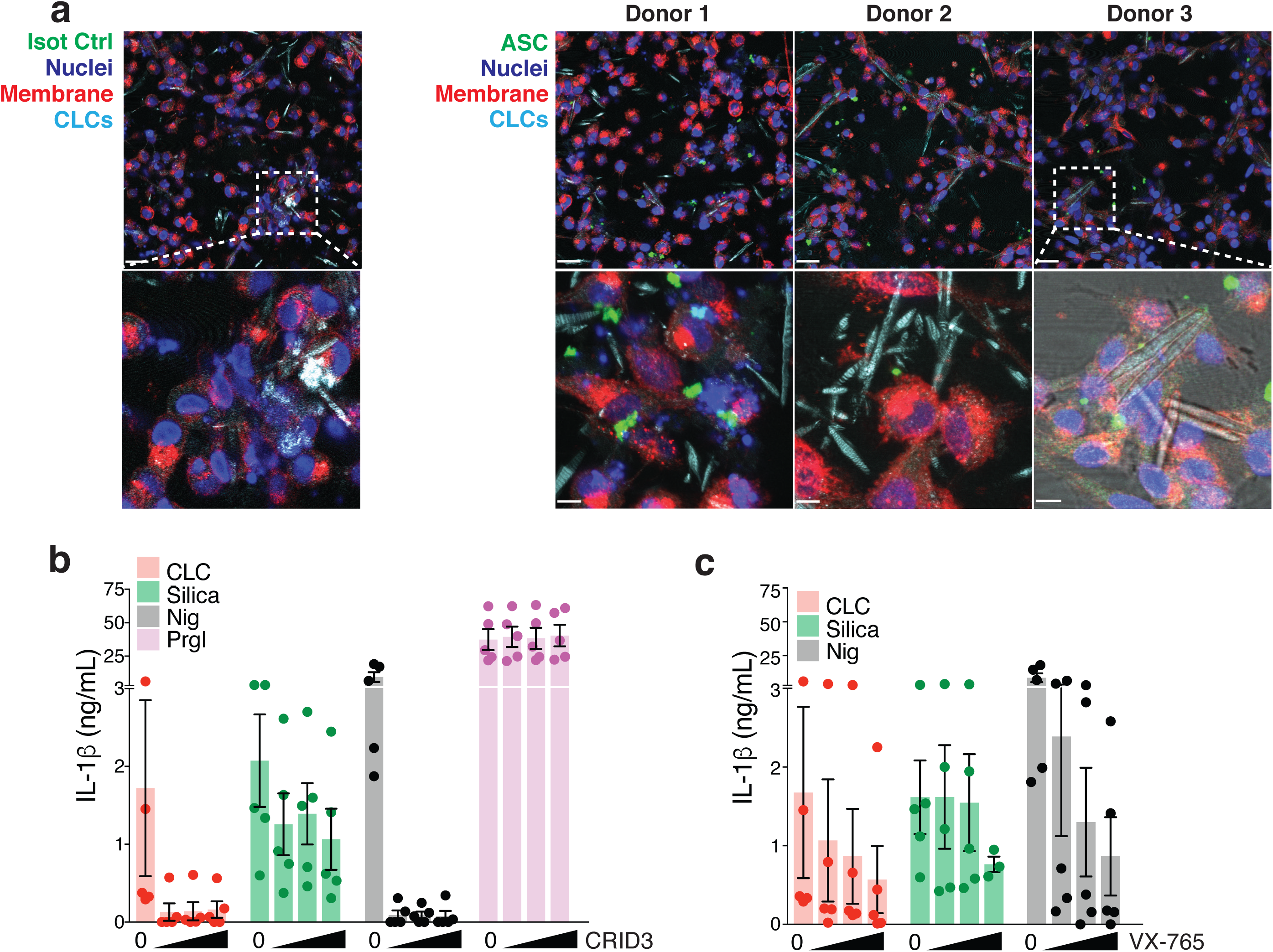
CLCs activate the NLRP3 inflammasome in human macrophages. (**a**) Confocal imaging of LPS-primed (2 ng/mL, for 3 hours) hMDMs stimulated with CLCs (2 per macrophage) for 6 hours. Cells were fixed and stained with directly Alexa Fluor 647 (AF647)-labeled anti-ASC antibody, or equal amounts of AF647-IgG isotype control mAbs. Plasma membrane (red, WGA-AF555), nucleus (deep blue, Hoechst 34580), ASC (green, AF647), and CLCs (light blue, laser reflection). Data is representative of two independent experiments. (**b**) HTRF measurement of IL-1β in LPS-primed (2 ng/mL, for 3 hours) human macrophages that were left untreated, or further stimulated with CLCs (2 per macrophage, 6 hours), silica crystals (100 μg/mL, 6 hours), cholesterol crystals (250 μg/mL, 6 hours), nigericin (10 μM, 1.5 hours), or PrgI (2 μg/mL, 3 hours) together with LFn-PA (0.5 μg/mL) in the presence of increasing concentrations of the NLRP3 inhibitor CRID3 (0, 2.5, 5, or 10 μM), or the caspase-1 inhibitor VX-765 (0, 5, 10, or 30 μM). Bars represent the mean and SEM of pooled data from five independent experiments. Each symbol represents values from a different donor. The same IL-1β data points for the CLC-treated hMDMs in the absence of inhibitors are represented in **b** and **c**.

In line with a role for the NLRP3 inflammasome in the intracellular sensing of CLCs, specific inhibition of NLRP3 and caspase-1 with the compounds CRID3 (Coll et al., 2015) and VX-765, respectively, ablated CLC-induced IL-1β release by hMDMs (Fig. 3b-c). In contrast, the response to NAIP-NLRC4 inflammasome activation, tested through the delivery of the needle protein of the *Salmonella* pathogenicity island 1 type III secretion system (PrgI) into the cytosol, was not affected by the NLRP3 inhibitor CRID3, demonstrating specificity. These findings indicate that, similar to silica (Hornung et al., 2008), uric acid (Martinon et al., 2006a), and cholesterol crystals (Duewell et al., 2010), CLCs induce IL-1β activation and release from hMDMs by activating the NLRP3 inflammasome.

### CLCs induce inflammasome activation *in vivo*

One of the main functions of IL-1 family cytokines is to induce the recruitment of immune cells, such as neutrophils and monocytes, which limits the spread of infection and initiates tissue repair (Dinarello, 2009). Inflammasome activation *in vivo* is characterized by strong neutrophil infiltration in tissues (Satoh et al., 2013), and tissue neutrophilia is a hallmark of IL-1-driven autoinflammatory diseases (Bonar et al., 2012; Broderick et al., 2015). We therefore used a well-established mouse peritonitis model (Yanagida et al., 2013; Duewell et al., 2010; Hornung et al., 2008) to assess whether CLCs also promote neutrophil infiltration, which could indicate that inflammasomes are activated *in vivo*. We injected wild-type C57BL/6 mice intraperitoneally with PBS or with CLCs. Silica crystals were used as a positive control, as these crystals were reported to cause IL-1R-dependent neutrophil infiltration *in vivo* (Hornung et al., 2008; Kuroda et al., 2011). CLCs caused peritonitis characterized by massive infiltration of neutrophils (CD11b^+^, Ly6G^+^, Ly6C^int^) and inflammatory monocytes (CD11b^+^, Ly6G^−^, Ly6C^+^) (Fig. 4a-b) into the peritoneal cavity. Of note, imaging of cells recovered from the PELF revealed the presence of neutrophils and monocytes that had phagocytosed CLCs (Fig. 4c and Fig. S2), confirming that CLCs are also phagocytosed *in vivo*.

**Fig. 4.**
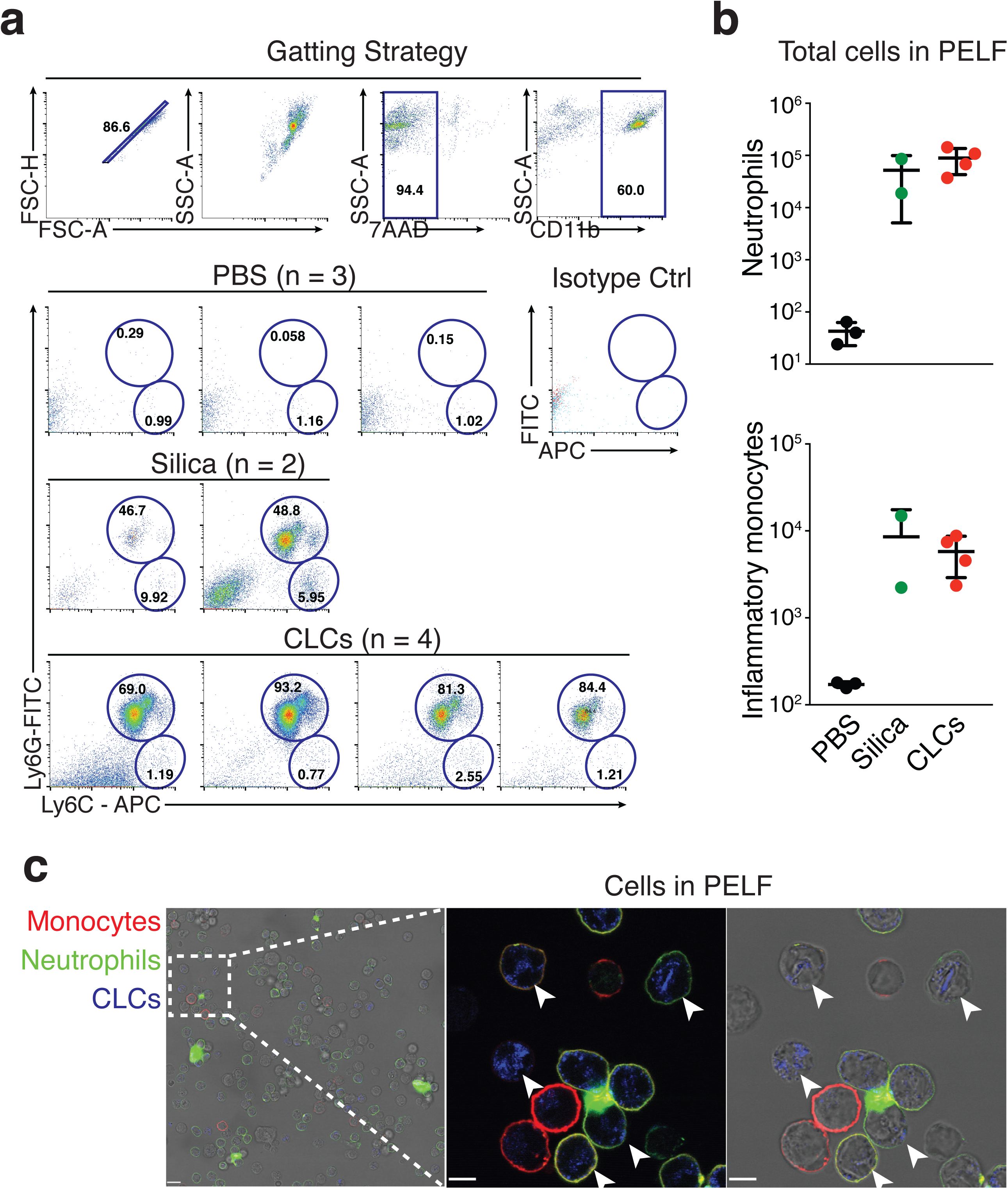
CLCs cause neutrophil recruitment *in vivo*. (**a**) Flow cytometric analysis showing gating strategy and dot-plots used for the quantification of cells in the peritoneal lavage fluids (PELF) from C57BL/6 mice 6 hours after intraperitoneal injection of 100 µL of PBS alone, or PBS containing 100 µg of silica crystals, or 1⨯10^6^ CLCs. Numbers in the regions show percentages of cells. (**b**) Flow cytometry quantification of live (7AAD^−^) neutrophils (CD11b^+^Ly6G^+^), and inflammatory monocytes (CD11b^+^Ly6C^+^Ly6G^−^) in the PELF from mice injected i.p. with PBS, or crystals. Each symbol represents an individual mouse; small horizontal lines indicate the mean and SD. (**c**) Confocal microscopy of the cells in the PELF of mice injected with CLCs as described in **b**. Monocytes (red, Ly6C-AF647), neutrophils (green, Ly6G-FITC), CLCs (blue, laser reflection). Arrowheads indicate CLCs, with their characteristic morphology, that were phagocytosed by neutrophils and monocytes. Scale bars: 42 μm left panel, 9 μm middle and right panels.

Although peritoneal injection of CLCs is useful to assess crystal-induced inflammation *in vivo*, it does not represent a physiologically relevant site for CLC formation. CLCs were frequently reported in the airways and sputum of patients with a variety of highly prevalent respiratory diseases, such as asthma (Dor et al., 1984). To confirm whether CLCs induce inflammasome activation *in vivo* in the lung and to locally associate CLCs uptake with inflammasome activation, we performed intratracheal instillation of CLCs into the lungs of ASC-mCitrine transgenic reporter mice (Tzeng et al., 2016). ASC-mCitrine mice were instilled with either PBS, CLCs, or silica crystals. Six hours after instillation, we assessed the infiltration of neutrophils, formation of mCitrine-fluorescent ASC specks, and production of pro-inflammatory cytokines in the bronchoalveolar lavage fluids (BALF). CLC-instilled mice displayed large numbers of neutrophils in their BALFs (Fig. 5a). Very few or no neutrophils (CD11b^+^, Ly6G^+^, Ly6C^int^) were found in the BALFs of mice instilled with PBS. In line with the observations from the peritonitis model, we found that cells recovered in BALF contained phagocytosed CLCs. This was accompanied with the formation of ASC-mCitrine specks, which was further validated with staining of ASC-mCitrine specks with an Alexa-647 directly conjugated anti-ASC antibody (Fig. 5b and S3). Measurements of IL1-β present in the BALFs showed a marked increase of this pro-inflammatory cytokine six hours after injection with silica crystals or CLCs, compared to PBS injected animals. The levels of TNF, an inflammasome-independent pro-inflammatory cytokine, were also elevated in the BALFs of mice treated with CLCs (Fig. 5c).

**Fig. 5.**
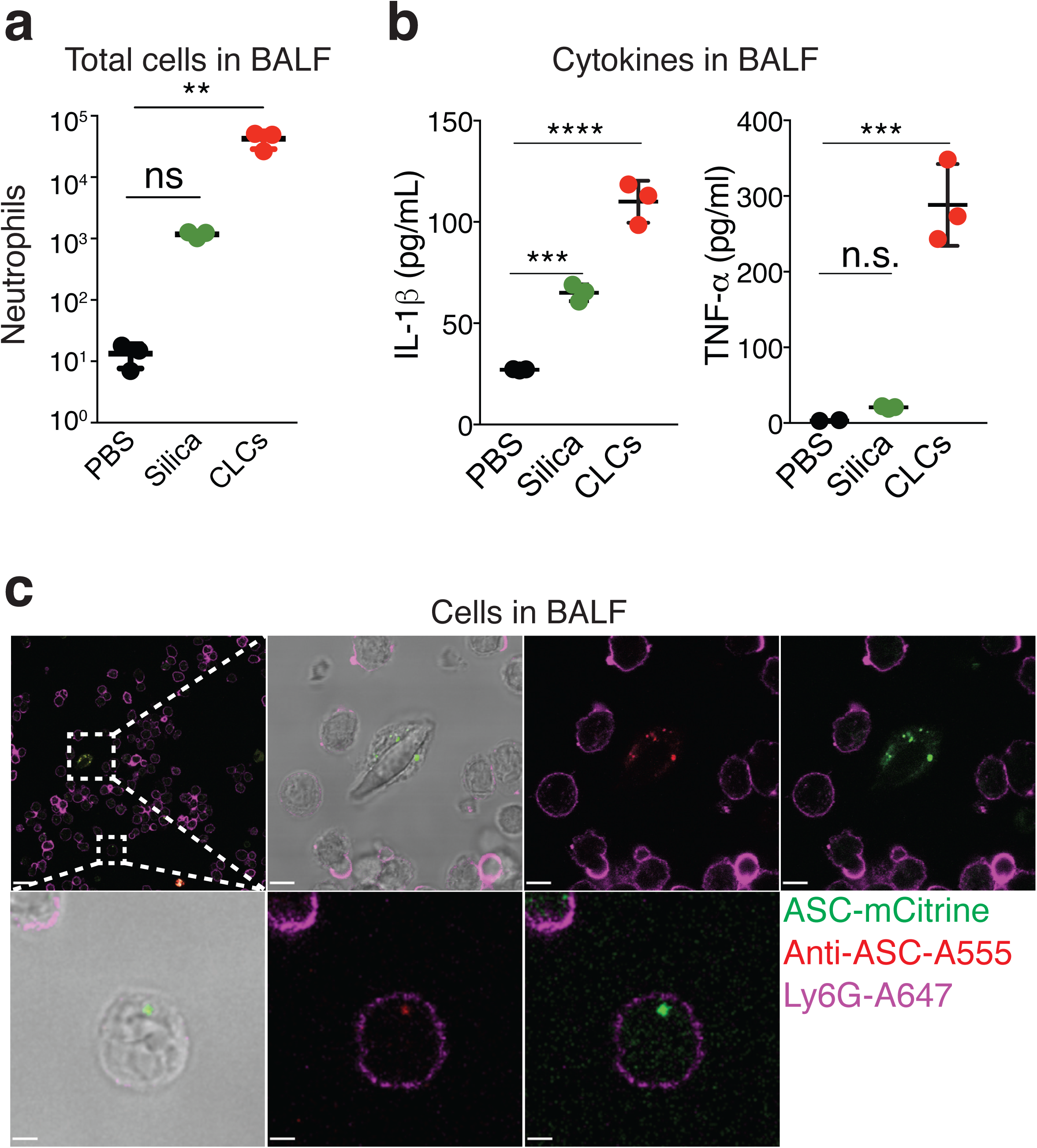
CLCs cause inflammasome activation in the lungs. (**a**) Flow cytometry quantification of live (7AAD^−^) neutrophils (CD11b^+^Ly6G^+^) in the bronchialveolar lavage fluid (BALF) of ASC-mCitrine transgenic mice 6 hours after intratracheal administration of 60 μL of PBS alone, or 60 μL of PBS containing 1 mg of silica crystals, or 3.5 × 10^6^ CLCs. Each symbol represents an individual mouse; small horizontal lines indicate the mean and SD. (**b**) Confocal imaging of cells present in the BALF of ASC-mCitrine transgenic mice instilled intratracheally with CLCs showing the formation of ASC aggregates in their cytosol. ASC (Green, mCitrine), Neutrophils (magenta, Ly6G-A647), anti-ASC (red, Anti-ASC A555. (**c**) IL-1β and TNF-α cytokine levels in the BALF of ASC-mCitrine transgenic mice treated as described in **a**. Each symbol represents an individual mouse; small horizontal lines indicate the mean and SD.

Altogether, our data show that CLCs induce inflammasome activation *in vivo* and suggest that their accumulation in tissues might be an important driver for chronic tissue inflammation following eosinophil infiltration.

### Concluding remarks

Our study identifies a previously unappreciated inflammatory capacity of CLCs and opens new avenues to consider the immunogenicity and potential relevance of these crystals for common chronic immunological diseases, such as asthma.

CLCs are found in the sputum of asthma patients with eosinophilia(Dor et al., 1984; Sakula, 1986). Although asthma patients can display heterogeneous phenotypes, an important molecular mechanism of asthma is type 2 inflammation, which is characterized by an increase in IgE titers, the recruitment of eosinophils, and the predominance of T_H_2 cells, which secrete IL-4, IL-5 and IL-13 (Fahy, 2015). Type 2 immunity can also influence the expression of classical pro-inflammatory cytokines, such as IL-1β, that can assist in the development of chronic inflammation. In line with these observations, endogenous danger associated molecular patterns (DAMPs), including uric acid crystals, traditionally known as the cause of inflammation in gout, are strong adjuvants for Th2 immunity (Kool et al., 2008; 2011).

Indeed, IL-1β levels have been shown to be increased in asthma, and the number of macrophages expressing IL-1β in the bronchial epithelium of asthma patients is higher in comparison with control subjects (Sousa et al., 1996). Our findings show that CLCs, which are commonly found in fluids and tissues of patients with eosinophilia, induce strong release of the highly pro-inflammatory cytokine IL-1β. This finding, together with the report of CLCs found in the phagosomes of macrophages *in vivo* (Lao et al., 1998; Dvorak et al., 1990), suggests that the uptake of CLCs by macrophages and the subsequent release of IL-1β can be one of the signals that perpetuate and exacerbate inflammation in the tissues of patients with eosinophilic disorders. On that note, IL-1β has been reported as a critical cytokine for the development of a Th2/Th17-predominant subtype in asthma patients (Liu et al., 2017). Th2/Th17-predominant asthma patients present dual-positive Th2/Th17 cells in their BALFs and exhibit a severe form of asthma. Moreover, these patients show increased eosinophil counts in their BALFs. Whether CLCs, which are a hallmark of eosinophilic diseases, are present in the BALFs of Th2/Th17-predominant asthma patients, and whether their recognition by alveolar macrophages are the cause of the increased levels of IL-1β in these patients is a matter of further research. In this scenario, inhibition of CLCs formation, engulfment, or effects in IL-1β release –potentially by blocking NLRP3 activity—could benefit patients with asthma.

The results presented in this study expand our current understanding on how protein crystallization contributes to immunogenicity and highlight the need to consider novel therapeutic strategies that neutralize protein crystallization lingering from the activity of granulocytes.

## ACKNOWLEGMENTS

We thank Meghan Sheehan, Cassandra C. Paul, and Michael A. Baumann, from the Wright State University (WSU) for the AML 14.3D10 cell line. We thank Maximillian Rothe for technical assistance. We thank Feng Shao (National Institute of Biological Sciences, Beijing 102206, China) for the plasmid encoding the PrgI-protective antigen conjugate protein, as well as Matthias Geyer and David Fußhöller for the purified PrgI protein used here to activate the NLRC4 inflammasome. B.S.F. is supported by a grant from the European Research Council (PLAT-IL-1) and the German Research Foundation (DFG, SFBTRR57). B.S.F, E.L. and N.G. are members of the ImmunoSensation cluster of Excellence. N.G. is supported by grants from the DFG (SFB704 and GRK2168). E.L. is supported by grants from the DFG (SFB645, 704, 670, 1123, TRR57, 83) and the European Research Council (InflammAct).

## AUTHOR CONTRIBUTIONS

E.L and B.D.F. conceived the study and B.D.F. and J.F.R.A. designed and performed experiments and analyzed the data. G.E. and C.S.M. provided technical support. M.A.A, W.K. and N.G. performed experiments and analyzed data. B.S.F. and J.F.R.A. wrote the manuscript with input from E.L.

## EXPERIMENTAL PROCEDURES

### Reagents

Ultrapure Lipopolysaccharide (LPS) from E. Coli 0111:B4 and nigericin were from Invitrogen, DRAQ5 was from eBiosciences. Cytochalasin D was purchased from Sigma and CA-074 Me from Enzo. The caspase-1 inhibitor Belnacasan (VX-765) was from Selleckchem. The NLRP3 specific inhibitor CRID3, (also known as Cytokine release inhibitory drug 3, or MCC950) was purchased from Sigma-Aldrich. HTRF kits for human IL-1β and TNF-α were from Cisbio Bioassays and were used according to manufacturer’s instructions. Cholesterol crystals were prepared as previously described(Zimmer et al., 2016). Briefly crystals were prepared from a 2 mg/ml cholesterol solution in 1-propanol. Crystallization was induced by addition of 1.5 volumes of endotoxin-free water. Crystals were dried and resuspended in sterile PBS.

### Mice

C57BL/6 mice were acquired from The Jackson Laboratory (Stock No. 000664). C57BL/6 ASC-mCtrine transgenic mice (B6.Cg-Gt(ROSA)26Sor^tm1.1(CAG-Pycard/mCitrine^*^,-CD2^*^)Dtg^/J) were acquired from The Jackson Laboratory (Stock No. 030744). Mice were housed in pathogen-free conditions and were handled in accordance with the Guide for the Care and Use of Laboratory Animals of the US National Institutes of Health and the Institutional Animal Care and Use Committee (IACUC) of the University of Massachusetts Medical School. C57BL/6 ASC-mCtrine transgenic mice (B6.Cg-Gt(ROSA)26Sor^tm1.1(CAG-Pycard/mCitrine^*^,-CD2^*^)Dtg^/J) were acquired from The Jackson Laboratory (Stock No. 030744).

### Cell lines

The AML 14.3D10 cell line was kindly gifted from Meghan Sheehan, Cassandra C. Paul, and Michael A. Baumann, from the Wright State University (WSU) Wright State University 3640 Colonel Glenn Highway, 304 University Hall, Dayton, OH 45435. AML14.3D10 cells were cultured in RPMI medium 1640 (Gibco) containing 10% fetal bovine serum (FBS) (Invitrogen), 1% penicillin/streptomycin, 1x GlutaMAX and 1x sodium pyruvate (both from Life Technologies) (complete medium).

### Human primary cells

Buffy coats from healthy donors were obtained according to protocols accepted by the institutional review board at the University of Bonn (local ethics votes Lfd. Nr. 075/14). Primary human macrophages were obtained through differentiation of CD14+ monocytes in a medium complemented with 500 U/mL rhGM-CSF (Immunotools) for 3 days. In brief, human peripheral blood mononuclear cells (PBMCs) were obtained from buffy coats of healthy donors by density gradient centrifugation in Ficoll-Paque PLUS (GE Healthcare). PBMCs were incubated at 4 °C with magnetic microbeads conjugated to monoclonal anti-human CD14 antibodies (Miltenyi Biotec). CD14+ monocytes were thereby magnetically labeled and isolated using a MACS column placed in a magnetic field as indicated by manufacturer (Miltenyi Biotec). Monocytes-derived macrophages were cultivated in complete medium during all experiments.

### Preparation of CLCs

CLCs were isolated following an adapted version of the protocol published elsewhere (Ackerman JI 1980). All steps were carried out at 4 °C to avoid protein degradation and under sterile conditions using a class II biosafety cabinet and sterile reagents. At least 600 × 10^6^ AML14.3D10 cells were washed twice with phosphate saline buffer (PBS) (Life Technologies) and resuspended in hypotonic (0.15% NaCl) PBS complemented with cOmplete protease inhibitor cocktail (Roche) (lysis buffer). For every mL of cell pellet, 1 mL of lysis buffer was used for resuspension of the cells. Resuspended cells were then lysed at 4 °C using a Dounce tissue grinder. 30 strokes were necessary for lysing >90% of the cells, according to inclusion of trypan blue stain. DNA and big membranes were removed by centrifugation of the lysates at 1,000 × g for 10 minutes at 4 °C. The pellets were resuspended in 500 μL of lysis buffer and the suspension was again centrifuged at 1,000 × g for 10 minutes at 4 °C to recover any soluble protein that could be trapped within the pellet of membranes. The supernatants from both centrifugation steps were combined and further centrifuged at 40,000 × g for 20 minutes at 4 °C to remove organelle membranes and cellular debris. The pellets were discarded and supernatants were centrifuged again at 40,000 × g for 30 minutes at 4 °C. The supernatants were combined and its volume was reduced to ~1mL by centrifugation at 3,200 × g in an Amicon Ultra-15 centrifugal filter with a membrane with NMWL of 10 kDa (EMD Millipore). This cleared and concentrated protein lysate already contained CLCs in suspension with rest of membranes and protein precipitates when it was observed under the microscope. CLCs were separated by centrifugation at 750 × g for 5 minutes at 4 °C. Supernatants were kept for 16 hours at 4 °C, and they were later on examined for the existence further CLCs. This second set of CLCs was separated by centrifugation at 750 × g for 5 minutes at 4 °C and mixed with the previous set. Combined CLCs were resuspended in cold PBS and thoroughly washed (a minimum of 6 times) until the CLCs were mostly free of contaminants when observing under the microscope. After all the washing steps, CLCs were counted and diluted with complete medium previous use for stimulation of macrophages.

### Protein gels and Western blot

Protein-enriched samples during the process of CLCs generation were mixed with NuPAGE® LDS Sample Buffer (4X) and with NuPAGE® Sample Reducing Agent (10X) (both from ThermoFischer Scientific) following to heating at 85 °C for 10 minutes to denaturalize and reduce all the proteins present in the samples.

Denaturalized and reduced proteins were separated in a NuPAGE 12% Bis-Tris protein gel (ThermoFischer Scientific) under electrophoretic conditions in MES buffer. After electrophoresis, proteins in the gel were stained with Coomassie Brilliant Blue R-250 dye (ThermoFischer Scientific) for ten minutes. Gels were destained in destaining solution (15% isopropanol + 10% acetone in water) for 16 hours. Alternatively, proteins in the gel were transferred into an Immobilon-FL PVDF membrane (Millipore). Nonspecific binding was blocked with 3% BSA in Tris-buffered saline (TBS) for 1 h, followed by overnight incubation with a rabbit monoclonal antibody directed against human Galectin-10 (EPR11197Abcam, dilution 1:10000) in 3% BSA in TBS with 0.1% Tween-20 (TBS-T). Membranes were washed three times in TBS-T and incubated for 2 hours with an IRDye 680RD Donkey anti-Rabbit antibody (Licor) in TBS-T. Membranes were washed two times with TBS-T and a last time with TBS before measuring fluorescent signal in an Odissey infrared imager model 9120.

### Immunocytochemistry

Cells were fixed in PBS containing 4% formaldehyde methanol free (Thermo Scientific) for 15 minutes at room temperature. After fixation, membranes were stained with wheat germ agglutinin (WGA)-Alexa Fluor 555 (Invitrogen) at 5 μg/mL in PBS for 10 minutes at room temperature. Fixed cells were incubated for 1 hour at room temperature with PBS containing 10% goat serum, 1% FBS, 0.5% Triton X-100 (blocking/permeabilising buffer), and then incubated overnight at 4 °C with 0.1 μg/mL of mouse anti-ASC monoclonal antibody (clone HASC-71, Biolegend) directly labeled with AF647 or its respective isotype control, also labeled with AF647. Cells were washed three times with blocking/permeabilising buffer. Nuclei were stained with DRAQ5 (1:5,000, Thermo Fischer).

### Microscopy

Confocal microscopy was combined with fluorescence microscopy using a Leica TCS SP5 SMD confocal system (Leica Microsystems, Wetzlar, Germany). Images were acquired using a 63X objective, with a numerical aperture of 1.2, and analysed using the Volocity 6.01 software (PerkinElmer, Waltham, Massachusetts, U.S.A.). For better visibility, in some instances, brightness and contrast were adjusted and all images within one experiment and applied equally to isotype controls and staining conditions.

### *In vivo* peritoneal injection of CLCs

C57BL/6 mice were exposed to crystals by the peritoneal injection of 100 uL of PBS containing either 1 × 10^6^ CLCs, 100 μg of silica crystals, or nothing. After 6 hours, mice were euthanized and 6 mL of lavage solution (RPMI + 3 mM EDTA) were injected into the peritoneum of mice as described elsewhere (Yanagida et al., 2013). After massaging the distended peritoneal cavity, the peritoneal lavage fluid (PELF) was recovered and separated into a cellular fraction and a liquid fraction by centrifugation at 350 × g at 4 °C during 5 minutes. These cells were assayed in flow cytometry to determine neutrophil recruitment and also observed under the microscope (Leica SP5) to identify cells that have phagocytosed CLCs.

### Intratracheal instillation of crystals

C57BL/6 ASC-mCitrine mice were instilled at a dose of 1 mg of silica crystals, or 3.5 × 10^6^ CLCs in 60 μL of PBS. The sample suspension was intratracheally instilled once to isoflurane anesthetized animals by a syringe through a catheter inserted into the airway. A control group was instilled with PBS alone. Mice were sacrified 6 hours after instillation, and 3 mL of bronchoalveolar lavage fluid was collected to determine the total cell count, neutrophil (7AAD^−^, CD11b^+^, Ly6G^+^) counts, and cytokine levels by HTRF.

### Neutrophil recruitment assays

Cells recovered in the PELF or BALF were resuspended in 2 mL of staining solution (PBS + 2% FCS) and counted with a hemocytometer. For neutrophils and monocytes identification, 250 uL of cells were stained with fluorochrome-labelled antibodies against CD11b (clone M1/70, Biolegend), Ly-6G (clone 1A8, BD Biosciences) and Ly-6C (clone ER-MP20, Bio-Rad), or their respective fluorochrome-labelled isotype Ig. After staining with 7AAD (BD Biosciences) for exclusion of dead cells, cells were analysed in a MACSQuant Analyzer 10 flow cytometer.

### Cytokine levels measurement in the BALF

The liquid fraction of the BALF was concentrated using Amicon Ultra-2 mL 10K centrifugal filters. The amounts of mouse IL-1β and mouse TNF-α in the concentrated BALF were quantified using a HTRF assay kit (Cisbio Bioassays) and normalized to the volume of concentrated lavage.

### Statistical analysis

Unless otherwise stated, all data is pooled from a minimum of 3 independent experiments carried out on macrophages from different donors. For clearance, individual experiments are depicted as symbols in graphs. Each symbol represents average values from technical triplicates from an individual donor. Data are graphed as mean, error bars show standard deviation (SD, when pooled from 2 independent experiments), or error of mean (SEM, when pooled from 3 or more independent experiments). The significance of differences between groups was evaluated by one-way analysis of variance (ANOVA) with repeated measures with Holm-Sidak’s postcomparison test and Geisser-Greenhouse correction. Statistical analysis was carried out with Graphpad Prism (version 6.0). Data were considered significant when p < 0.05 (*), 0.01 (**), 0.001(***), or 0.0001(****).

## Supplementary Figures

**TABLE S1:** Source data for all the figure panels with human pooled IL-1β data from different donors.

**Fig. S1:** Detailed schematics of CLC production from AML14.3D10 cells with protein characterization through all washing steps involved. 30 μg of the resultant AML14.3D10 lysates after each centrifugation step (see Experimental Procedures), 3 μg of pure CLCs were loaded into a protein gel.

**Fig. S2:** Confocal microscopy of the cells in the PELF of wild-type C57BL/6 mice 6 hours after intraperitoneal injection of 100 μL of PBS alone, or PBS containing 1×10^6^ CLCs. Note that for the CLC-injected mice, cells show phagocytosed CLCs.

**Fig. S3:** Confocal imaging of cells present in the BALF of ASC-mCitrine mice instilled intratracheally with CLCs or silica crystals showing the formation of ASC aggregates in their cytosol. Note that ASC remains mainly evenly distributed in the cytosol of the cells exposed to only PBS. ASC (Green, mCitrine), Neutrophils (magenta, Ly6G-A647), anti-ASC (red, Anti-ASC A555.

**Movie S1:** Time-lapse confocal imaging of human macrophages phagocytosing CLCs.

